# Artificial sweeteners differentially activate sweet and bitter gustatory neurons in *Drosophila*

**DOI:** 10.1101/2025.02.06.636883

**Authors:** Christian Arntsen, Jake Grenon, Isabelle Chauvel, Stéphane Fraichard, Stéphane Dupas, Jérôme Cortot, Kayla Audette, Pierre-Yves Musso, Molly Stanley

## Abstract

Artificial sweeteners are highly sweet, non-nutritive compounds that have become increasingly popular over recent decades despite research suggesting that their consumption has unintended consequences. Specifically, there is evidence suggesting that some of these chemicals interact with bitter taste receptors, implying that sweeteners likely generate complex chemosensory signals. Here, we report the basic sensory characteristics of sweeteners in *Drosophila*, a common model system used to study the impacts of diet, and find that all noncaloric sweeteners inhibited appetitive feeding responses at higher concentrations. At a cellular level, we found that sucralose and rebaudioside A co-activated sweet and bitter gustatory receptor neurons (GRNs), two populations that reciprocally impact feeding behavior, while aspartame only activated bitter cells. We assessed the behavioral impacts of sweet and bitter co-activation and found that low concentrations of sucralose signal appetitive feeding while high concentrations signal feeding aversion. Finally, silencing bitter GRNs reduced the aversive signal elicited by high concentrations of sucralose and significantly increased sucralose feeding behaviors. Together, we conclude that artificial sweeteners generate a gustatory signal that is more complex than “sweetness” alone, and this bitter co-activation has behaviorally relevant effects on feeding that may help flies flexibly respond to these unique compounds.

## INTRODUCTION

Artificial sweeteners are intensely sweet, non-nutritive compounds that have emerged as popular way to reduce caloric intake while maintaining palatability^1^. Sweeteners have become an increasingly common feature of modern diets^2^, a trend that is expected to persist^3^. The FDA and other regulatory agencies have currently approved several artificial sweeteners for general consumption, including sucralose, saccharin, aspartame, neotame, advantame, acesulfame potassium, and sugar alcohols^4,5^. Beyond these synthetic chemicals, naturally derived compounds like stevia and its extracts, stevioside and rebaudioside A (Reb A), have also grown in popularity as low-calorie sweet additives^6^. While these sweeteners are considered biologically inert and safe for consumption^7^, their increased prevalence has led to apprehension about their long-term safety^8^. The World Health Organization (WHO) has recently advised against using sweeteners for weight loss or dieting, citing potentially harmful physiological effects^9^. Both animal and clinical investigations have suggested gastrointestinal^10–13^, metabolic^14–16^, and cardiovascular^5,17,18^ risks associated with sweetener consumption. Many of these concerns are related to the unique sensory attributes of these chemicals, which can be hundreds to thousands of times sweeter than sucrose^19^, and the potentially harmful implications of experiencing high-intensity sweetness dissociated from energy intake^14^. Additionally, artificial sweeteners are detected by human sweet taste receptors (T1R2/T1R3)^20–22^, but *in vitro* evidence suggests that sucralose also binds to bitter receptors (T2Rs)^23,24^, including those expressed extra-orally^25^. These findings align with behavioral evidence indicating that sucralose and other sweeteners possess a bitter “aftertaste” quality for both humans and rodents^26–28^. However, the exact cellular mechanisms and behavioral consequences of sweetener chemosensation remain unclear, particularly regarding potential off-target bitter signaling.

*Drosophila melanogaster* has been an advantageous model organism for studying both gustatory processing^29^ and how artificial sweeteners impact multiple phenotypes^30–41^, including feeding. Initial investigations into sweetener feeding behavior demonstrated that flies preferentially consume, sucralose, aspartame, and saccharin, implying a conserved attraction to these compounds^42^. Later research examined the long-term effects of sweetener exposure and proposed that dietary sucralose leads to enhanced sweet taste sensitivity and increased food intake via a neuronal starvation response^39^. In contrast, another study found that sucralose deters feeding^41^, leading to further controversy about the effects of dietary sweeteners^40^. However this debate focused on feeding phenotypes that did not consider how sucralose is actually detected at a sensory level.

In flies, taste processing is initiated by peripheral gustatory receptor neurons (GRNs). GRNs are distributed throughout several structures, but most prominently within taste sensilla located on the fly’s primary mouthpart - the labellum^43,44^. Labellar GRNs detect chemical stimuli and directly transmit taste signals to a brain region known as the subesophageal zone (SEZ) for higher-order processing^45^. Among the five recognized GRN classes, sugar-sensing “sweet” cells are the primary appetitive GRN population and express sugar-specific gustatory receptors, such as *Gr64f*^46–48^. In contrast, bitter GRNs represent the main aversive GRN population and detect unpalatable compounds via several types of taste receptors, including *Gr66a*^45,49^. Accordingly, activation of sweet or bitter GRNs has strong but opposite effects on feeding^49,50^. Taking advantage of the abundant genetic and neurobiological tools available in *Drosophila*^51,52^, previous investigation of these taste cell populations has revealed complex and combinatorial mechanisms of gustatory processing for several classes of tastants^48,53–57^.

While prior research has indicated mild attraction to sweeteners at a behavioral level^42^, they only looked at the effects of stimulating taste cells in the legs (tarsal GRNs). How flies respond to sweetener stimulation on the labellum, where the majority of GRNs are located^58^, remains unclear. Moreover, the cellular responses elicited by these compounds in *Drosophila* have not been described. Here, we recorded behavioral responses to labellar sweetener stimulation, showing that flies exhibit concentration-dependent taste responses to different compounds. We then characterized the neuronal responses to sucralose and found that it co-activates sweet and bitter GRNs to modulate feeding behavior. We also included aspartame and Reb A in these analyses to demonstrate that bitter cellular responses extend to sweeteners with different chemical structures. Together, our findings shed light on the basic chemosensory underpinnings of artificial sweeteners and specifically highlight bitter taste cell co-activation as an important consideration for future sweetener research.

## RESULTS

### Flies exhibit dose-dependent taste responses to multiple artificial sweeteners

We first sought to describe the taste responses to artificial sweeteners at a behavioral level. To accomplish this, we assembled a panel of common sweeteners and tested how flies respond to each compound using the proboscis extension response (PER) assay^59^, specifically stimulating a fly’s labellum with a test solution and recording if they extend their proboscis to initiate feeding (Fig. 1A). The proportion of flies that extend to each test stimulation can be quantified and used as a measure of appetitive taste responsiveness. We performed PER experiments using a concentration series for each of the seven sweeteners in our panel: sucralose, saccharin, cyclamate, aspartame, neotame, rebaudioside A (Reb A), and maltitol. Across our panel, we observed dose-dependent taste responses to the different sweeteners, with most compounds eliciting the highest level of PER at low/moderate concentrations (Fig. 1).

**Figure 1:**
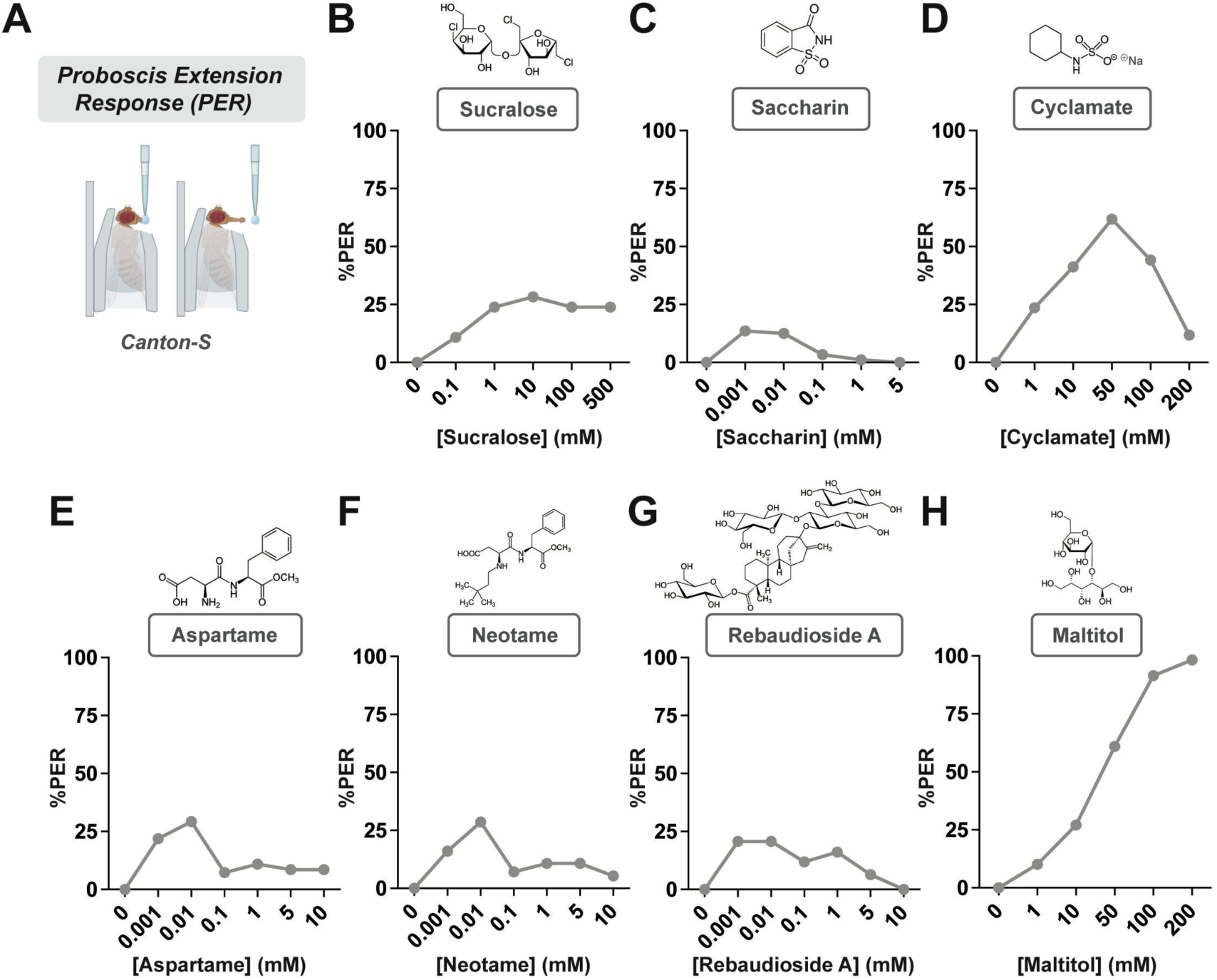
Flies exhibit diverse and concentration-dependent taste responses to a panel of artificial sweeteners. (A) Labellar proboscis extension response (PER) assay schematic showing stimulation with a test solution and a resulting extension. (B-H) Labellar PER with a panel of seven sweeteners presented to flies in a concentration series. %PER refers to the fraction of flies that extended to each respective concentration. *n* = 46-99. All data from mated female *Canton-S* flies.

Stimulating with cyclamate, a relatively mild sweetener used widely in Europe^60^ resulted in higher PER rates compared to the other sweeteners (Fig. 1D). However, cyclamate is the sodium salt of cyclamic acid, and its PER curve resembles that of sodium^48,61^, suggesting these flies likely detect and respond to the salt taste in addition to any of cyclamate’s sweetness.

Unlike the other non-nutritive compounds in our panel, maltitol is a semi-caloric sugar alcohol with a similar level of sweetness as sucrose in humans^62^. Maltitol exhibited the strongest responses of any sweetener we tested and PER rates increased at higher concentrations (Fig. 1H), mimicking sucrose PER^63,64^ and implying that flies are more strongly attracted to even partially nutritive sweeteners compared to non-caloric alternatives. Responses to saccharin, aspartame, neotame, and Reb A all shared a similar pattern - a small peak at 0.001 - 0.01 mM followed by very minimal responses at higher concentrations (Fig. 1C, 1E, 1F, 1G). In comparison, flies were more attracted to sucralose, with responses peaking around 10 mM and remaining consistent at higher concentrations (Fig. 1B). Overall, most of these high-intensity sweeteners elicited only mild attraction, indicating that flies may be responding to additional gustatory signals beyond just saccharinity.

### Sucralose activates sweet taste cells to promote feeding attraction

To more thoroughly characterize these taste responses, we explored how artificial sweeteners are encoded by the fly gustatory system at a cellular level. We first focused on sucralose because of its relatively strong taste responses (Fig. 1B) and its prominence in previous fly sweetener research^37–42^. As sucralose is considered 600 times sweeter than sucrose for mammals^65^, we began by analyzing the neuronal responses to sucralose detection in sweet GRNs (*Gr64f*+), the primary appetitive taste cell population in *Drosophila*. This functional analysis was accomplished using *in vivo* calcium imaging of labellar GRNs. In these experiments, we stimulated flies’ labella with a range of sucralose concentrations and recorded changes in calcium fluorescence from GRN axon terminals in the SEZ via selective *GCaMP6f* expression. This technique has been previously used in *Drosophila* to uncover nuances in gustatory encoding and processing^48,53,54,66–69^. We found that stimulation with 100 and 500 mM sucralose elicited significant increases in sweet GRN calcium activity compared to water, a negative control (Fig. 2A), indicating that these appetitive taste cells do respond to this sweetener. Interestingly, the magnitude of these responses was surprisingly moderate, particularly when compared to the 1 M sucrose positive control response level (Fig. 2A).

**Figure 2:**
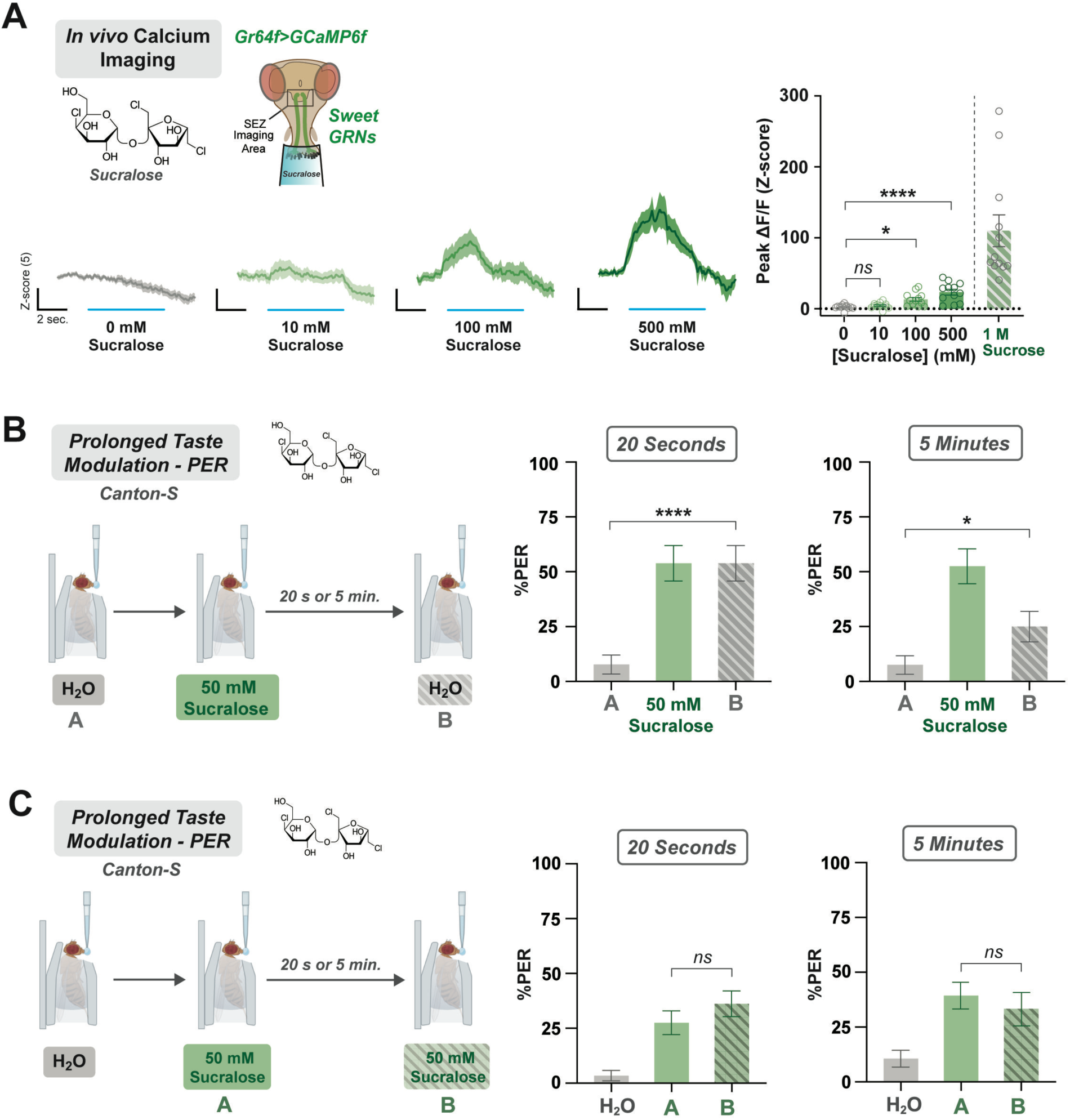
Sucralose activates sweet GRNs to signal feeding attraction at low concentrations. (A) *In vivo* calcium imaging of sweet GRNs (*Gr64f>GCaMP6f*) during labellar sucralose stimulation. Calcium responses measured as ΔF/F (Z-score) over time at each concentration (left) and peak ΔF/F (right). Blue lines under each curve indicate when the stimulus was on the labellum. *n* = 12 flies. (B) Prolonged taste modulation PER paradigm consisting of three consecutive stimulations: an initial water stimulation A, a 50 mM sucralose stimulation, and a delayed water stimulation B. Modulation assessed after 20 seconds (right) or 5 minutes (left). *n* = 40 flies for both experiments. All data from *Canton-S* flies. (C) Prolonged taste modulation PER paradigm with 50 mM sucralose stimulation A followed by a second, delayed 50 mM sucralose for stimulation B. *n* = 66-69 flies. All data from *Canton-S* flies. All experiments used mated females. Data plotted as mean ± SEM. *ns =* no significance, *\*p < 0.05*, *\*\*\*\*p < 0.0001* by ordinary one-way ANOVA with Dunnett’s multiple comparisons test (A) or Wilcoxon matched-pairs signed rank test (B, C).

We then asked if this level of sweet activation represents a signal strong enough to impact feeding. Previous research on sweet GRNs revealed that both sucrose stimulation and optogenetic activation of these cells were sufficient to enhance subsequent PER to neutral stimuli, even after a prolonged delay^50^. This finding suggests that activation of sweet GRNs primes the fly to respond appetitively to upcoming gustatory stimuli, thereby encouraging feeding. To determine if sucralose also impacts feeding in this manner, we utilized a similar prolonged taste modulation PER paradigm. Flies were presented with three consecutive stimulations: a baseline neutral water stimulation, a sucralose stimulation, and a second, delayed water stimulation (Fig 2B). We found that stimulating flies with 50 mM sucralose, a relatively low and attractive concentration (Fig. 1B), significantly enhanced PER to the second water stimulus both acutely (20 seconds) and after a prolonged delay (5 minutes) (Fig. 2B).

Next, we tested whether exposure to sucralose would also enhance the responses to a second 50 mM sucralose stimulation, but there was no change (Fig. 2C). This difference implies that the appetitive signal elicited by 50 mM sucralose specifically enhances taste responses to neutral stimuli and not for compounds that are already attractive. Together, these findings suggest that sucralose activates sweet GRNs and that a relatively low concentration signals feeding attraction up to 5 minutes after detection.

### Sucralose activates bitter taste cells to limit feeding

While higher concentrations of sucralose produced increasing calcium responses in sweet GRNs (Fig. 2A), our PER experiment instead showed that behavioral responses plateaued at these same concentrations (Fig. 1B). We reasoned that the discrepancy between our behavioral and cellular results could be explained by additional GRN populations responding to sucralose and forming a more complex gustatory signal. We hypothesized that aversive GRNs also respond to sucralose stimulation and that the resulting signal has biologically relevant effects on feeding. Using the same *in vivo* calcium imaging approach, we assessed how bitter GRNs (*Gr66a+*), the primary aversive GRN population, respond to sucralose detection. We found that stimulation with 100 and 500 mM sucralose significantly activated bitter GRNs (Fig. 3A). Unlike the mild calcium signals seen in sweet GRNs (Fig. 2A), the levels of bitter GRN activation by sucralose were much higher and comparable in magnitude to those elicted by the caffeine positive control stimulation (Fig. 3A).

**Figure 3:**
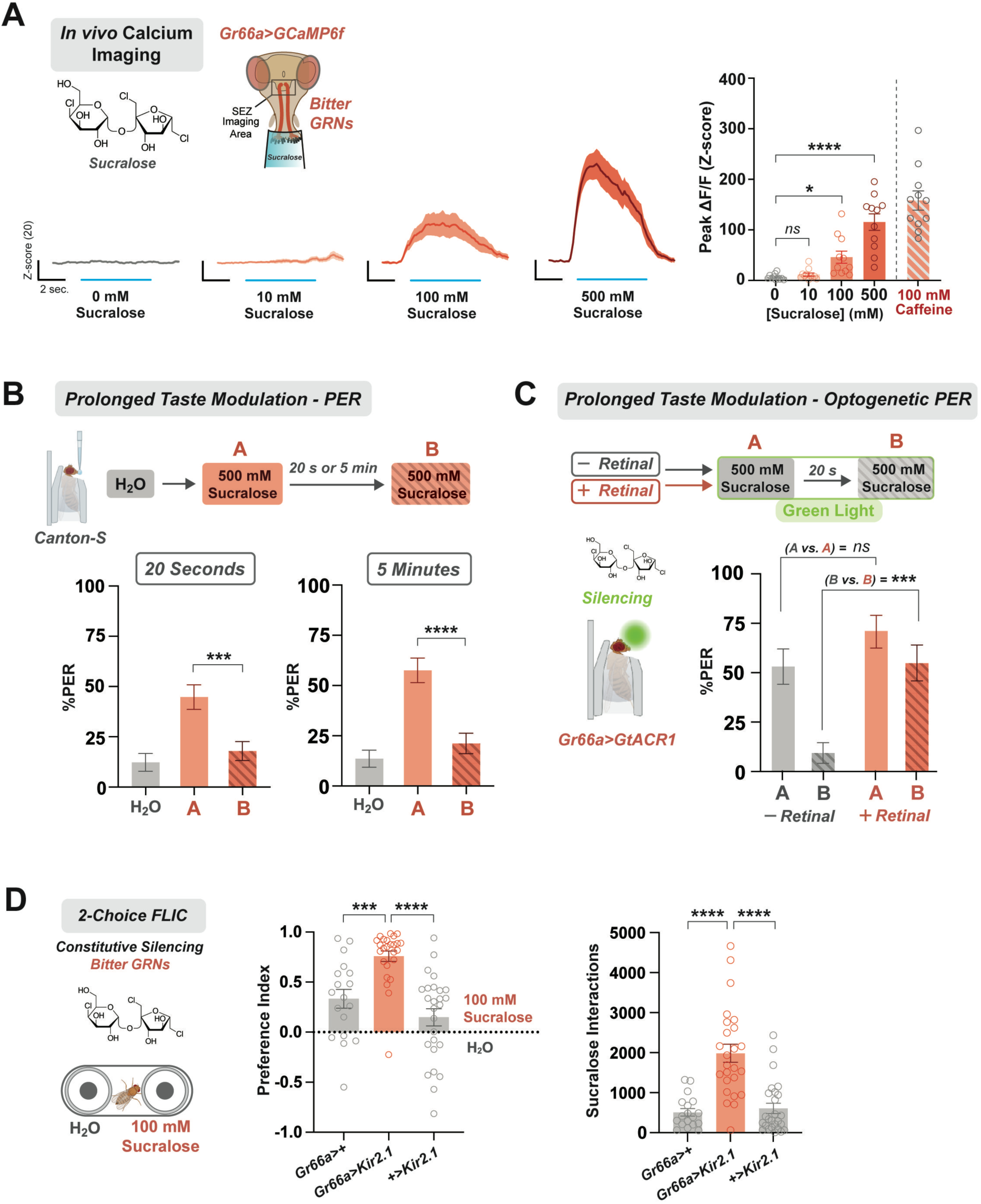
Sucralose activates bitter GRNs to signal feeding aversion at high concentrations. (A) *In vivo* calcium imaging of bitter GRNs (*Gr66a>GCaMP6f*) during labellar sucralose stimulation. Calcium responses measured as ΔF/F (Z-score) over time at each concentration (left) and peak ΔF/F (right). Blue lines under each curve indicate when the stimulus was on the labellum. *n* = 11 flies. (B) Prolonged taste modulation PER paradigm consisting of two repeated 500 mM sucralose stimulations separated by 20 seconds or 5 minutes. *n* = 66-67 flies. All data from *Canton-S* flies. (C) Optogenetic silencing of bitter GRNs (*Gr66a>GtACR1*) during prolonged taste modulation PER assay. Flies received two 500 mM sucralose stimulations separated by 20 seconds while exposed to green light. %PER to the initial A and delayed B stimulations were compared between flies pre-fed retinal (+Retinal) and control flies (-Retinal). *n* = 32 flies per condition. (D) Constitutive silencing of bitter GRNs (*Gr66a>Kir2.1*) during two-choice FLIC feeding assay. Flies were allowed to choose between two food sources (H_2_O and 100 mM sucralose) for three hours. Preference index (left) calculated based off number of interactions with each food option, sucralose interactions (right). *n* = 18-26 flies per genotype. All experiments used mated females. Data plotted as mean ± SEM. *ns =* no significance, *\*p < 0.05*, *\*\*\*p < 0.001, ****p < 0.0001* by ordinary one-way ANOVA with Dunnett’s multiple comparisons test (A, D) or Wilcoxon matched-pairs signed rank test (B, C).

Next, we tested if bitter GRN activation by sucralose represented a salient aversive feeding signal. Since 500 mM sucralose exhibited the strongest bitter GRN activation (Fig. 3A), we used this concentration as the representative bitter stimulus in our prolonged taste modulation PER assay. It was previously established that this paradigm also detects the ability of bitter GRN activation to suppress future taste responses^50^. We tested if initial exposure to 500 mM sucralose could produce a bitter signal strong enough to attenuate subsequent taste responses to the same 500 mM stimulation. Flies responded strongly to the initial sucralose stimulation, but PER to the second was significantly weaker after either a 20-second or 5-minute delay (Fig. 3B). The reduced responses to the second stimulation suggest the initial detection produced an aversive “aftertaste” signal that diminished sucralose attraction.

To validate that this sucralose-induced aversive signal is driven by bitter GRN activation, we predicted that silencing these cells would dampen the aversive component of sucralose and enhance taste responses as a result. We optogenetically silenced bitter GRNs via selective expression of *GtACR1*, an anion channel that hyperpolarizes neurons upon green light stimulation^70^. This approach also requires pre-feeding with all-*trans*-retinal, a chromophore necessary for proper GtACR1 function^66,71,72^. *Gr66a>GtACR1* flies without retinal pre-feeding served as experimental controls. We implemented this optogenetic manipulation into our aversive prolonged taste modulation PER assay by exposing flies to green light during two 500 mM sucralose stimulations separated by a 20-second delay (Fig. 3C). Consistent with our prediction, taste responses to the initial stimulation were similarly strong for both groups, but the flies pre-fed with retinal exhibited a significantly higher level of PER to the second stimulation compared to controls (Fig. 3C). These findings suggest that silencing bitter GRNs reduces the aversive signal elicited by 500 mM sucralose, confirming that the activation of these neurons drives the unpalatable aspect of sucralose detection.

To further assess the impact of bitter GRN activity on prolonged sucralose feeding, we used the Fly Liquid Interaction Counter (FLIC) to quantify nuanced feeding metrics^68,73^. For this assay, flies were individually loaded into feeding chambers containing two food options: water and 100 mM sucralose (Two-choice FLIC). Over a three-hour period, preference index, interactions, feeding events, and mean event duration were recorded for each fly. We tested how these sucralose feeding metrics changed when bitter GRNs were genetically inactivated via Kir2.1, an inward-rectifying potassium channel that constitutively silences neurons^74^. Flies with silenced bitter GRNs showed increased preference, interactions, and events with sucralose compared to genetic controls (Fig. 3D, Fig. S1A). To confirm these results with acute silencing, we repeated our 2-choice FLIC experiment in flies with bitter GRNs optogenetically silenced by green light exposure throughout the assay^75^. Compared to controls, flies pre-fed with retinal exhibited a stronger sucralose preference, but there were no differences in the other metrics (Fig. S1B). Collectively, our results demonstrate that sucralose co-activates sweet and bitter GRNs to form a concentration-sensitive gustatory code comprised of opposing attractive and aversive signals that reciprocally modulate feeding.

### Aspartame and Reb A minimally activate GRNs to impact feeding behaviors

To broaden our investigation of sweetener chemosensation, we assessed the cellular and behavioral responses for two additional chemicals: aspartame and Reb A. We chose these two compounds based on their popularity^76,77^ and to ensure our analysis included sweeteners with diverse chemical structures. In our initial PER experiments, both aspartame and Reb A produced minimal taste responses, particularly at high concentrations (Figs. 1E, 1G). If the fly gustatory system encodes these chemicals like sucralose, this response pattern could indicate sweet and bitter GRN co-activation. Surprisingly, we did not observe any cellular responses to aspartame in sweet GRNs (Fig. 4A, S2A). However, aspartame stimulation induced significant calcium activity in bitter cells (Figs. 4B, S2B), including an OFF peak with stimulus removal, a feature that has been observed with other aversive tastants^78,79^. Despite clear bitter activation, optogenetic silencing of bitter GRNs during low or high concentration aspartame stimulation had no effect on taste responses in our prolonged taste modulation PER assay (Fig S2C). Moreover, constitutive bitter GRN silencing did not alter aspartame feeding behavior (Fig. 4C, S2D), implying either that this signal is not salient for feeding or that without a simultaneous appetitive signal, aspartame will never be preferred.

**Figure 4:**
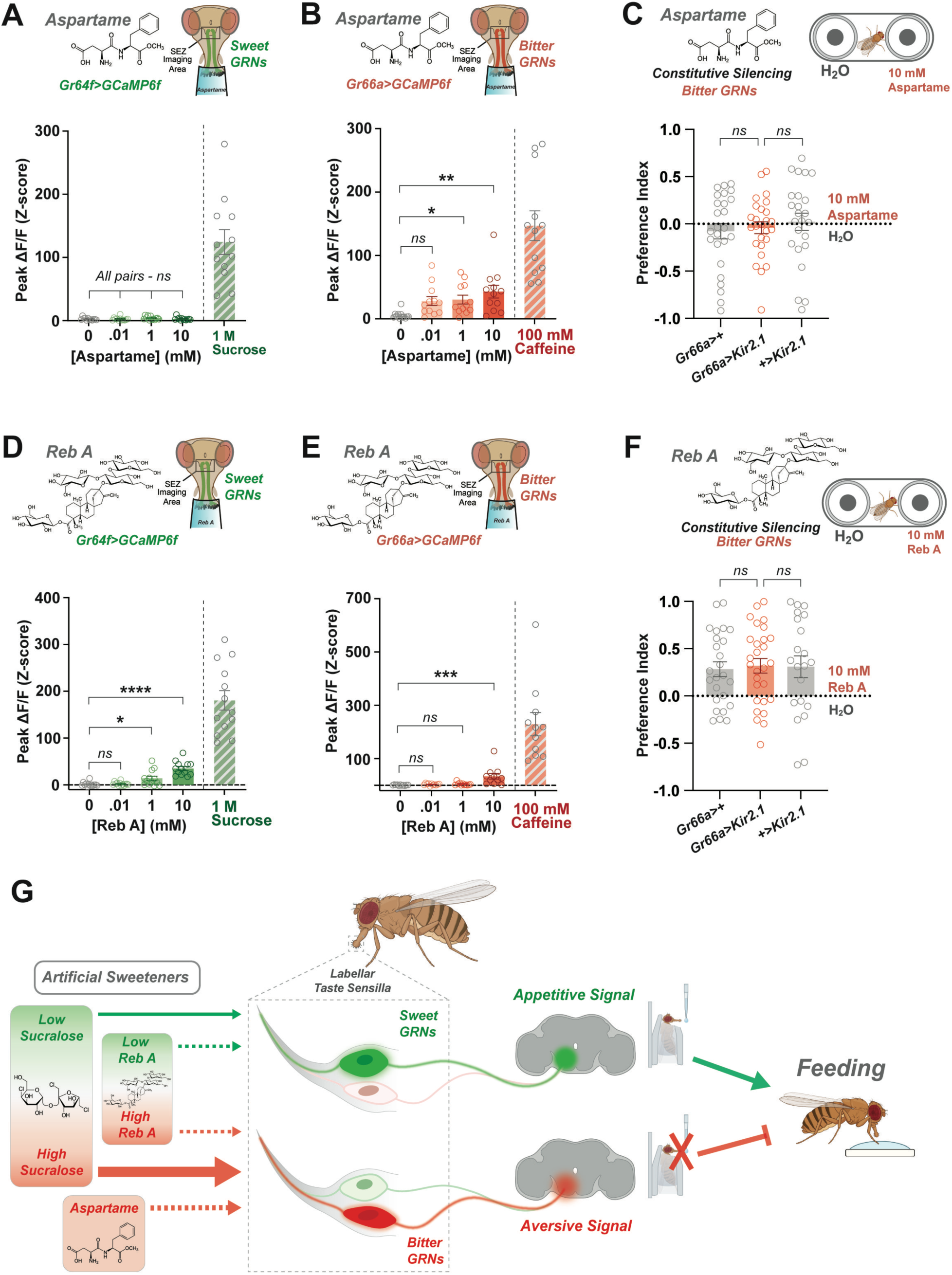
Aspartame and Reb A exhibit limited GRN responses that minimally impact feeding. (A, B) *In vivo* calcium imaging of sweet (A) and bitter (B) GRNs during labellar aspartame stimulation. Calcium responses measured as peak ΔF/F (Z-score). *n* = 12 flies per experiment. (C) Constitutive silencing of bitter GRNs (*Gr66a>Kir2.1*) during two-choice FLIC feeding assay. Flies were allowed to choose between two food sources (H_2_O vs. 10 mM aspartame) for three hours. Preference index calculated based off number of interactions with each food option. *n* = 26-27 flies per genotype. (E, F) *In vivo* calcium imaging of sweet (E) and bitter (F) GRNs during labellar Reb A stimulation. Calcium responses measured as peak ΔF/F (Z-score). *n* = 13 (E) and 11 (F). (G) Constitutive silencing of bitter GRNs (*Gr66a>Kir2.1*) during two-choice FLIC feeding assay. Flies were allowed to choose between two food sources (H_2_O vs. 10 mM Reb A) for three hours. Preference index calculated based off number of interactions with each food option. *n* = 21-28 flies per genotype. (H) Proposed model of how sweet and bitter GRNs encode several artificial sweeteners and how these signals impact feeding. Red-green gradients show how low concentrations of sucralose and Reb A predominantly activate sweet GRNs while higher concentrations co-activate bitter GRNs. Aspartame only activates bitter GRNs. Solid lines represent signals that affect feeding while dotted lines represent signals that do not significantly impact feeding. All experiments used mated females. Data plotted as mean ± SEM. *ns =* no significance, *\*p < 0.05*, *\*\*p < 0.01, ***p < 0.001, ****p < 0.0001* by ordinary one-way ANOVA with Dunnett’s multiple comparisons test.

In contrast, Reb A mildly activated both sweet and bitter GRNs. Sweet GRNs exhibited increased calcium activity when stimulated with Reb A (Figs. 4D, S3A) similar to sucralose (Fig. 2A). Additionally, the magnitude of Reb A’s bitter responses was relatively low compared to the other sweeteners (Figs. 4E, S3B). Silencing bitter GRNs had no impact on Reb A taste response modulation (Fig. S3C) or Reb A feeding, which did show a mild positive preference (Fig. 4F, S3D). The lack of behavioral phenotypes following bitter silencing implies that the aversive signals are insufficient to override the appetitive signals. Overall, we propose that multiple artificial sweeteners are sensed by both appetitive and aversive GRN populations (Fig. 4G). Sweet GRNs mediate appetitive responses to low concentrations of sucralose and Reb A, while bitter GRNs respond to aspartame and higher concentrations of sucralose and Reb A. Sweet and bitter co-activation by sucralose generates a balanced gustatory signal that modulates feeding in a concentration-dependent manner. In contrast, the distinct GRN responses to aspartame and Reb A have minimal effects on feeding behavior (Fig. 4G).

## DISCUSSION

Mammalian research into artificial sweeteners has focused on providing evidence for any potentially harmful effects with long-term sweetener intake^5,8,10–14,16–18^, but the mechanisms underlying pathological risks remain unclear, as does our understanding of how sensory systems signal the presence of sweeteners to the brain. Despite evidence indicating that bitter aftertastes associated with sweeteners are potentially due to bitter receptor (T2R) binding^23,24^, characterizing the impacts of sweetener sensation or consumption requires *in vivo* analyses that are currently challenging to achieve with mammals. These limitations have helped *Drosophila* become a common model organism for studying how dietary sweetener exposure impacts lifespan, feeding, and other phenotypes^30–42^. However, these studies may have limited relevance for mammals and humans if the sensory experience associated with these compounds differ between species. In the current study, we described the cellular basis of *Drosophila* sweetener gustation *in vivo* and explored how bitter taste cell co-activation impacts sweetener feeding behavior.

### Artificial sweeteners induce responses in both sweet and bitter gustatory neurons

Our labellar PER experiments showed only mild sweetener attraction – implying that flies may be experiencing more complex taste signals. To confirm, we recorded calcium responses from sweet and bitter GRNs during *in vivo* sweetener stimulation. This technique has successfully been used to describe mechanisms of peripheral taste processing in living, awake flies^48,53,54,66–69^, an approach that is currently not feasible with mammalian model systems. Sweet GRNs responded to both sucralose and Reb A, but not to aspartame. Bitter GRNs responded strongly to sucralose and moderately to aspartame and Reb A (Fig. 4G). These results indicate that sucralose may represent a well-conserved sweet ligand for human and *Drosophila* taste receptors, potentially due to its structural similarities with sucrose. However, the molecular basis of sucralose detection remains unknown. *In vitro* studies with mammalian cells has helped reveal that two bitter receptors expressed in human taste papillae, TAS2R43 and TAS2R44, specifically detect saccharin and acesulfame K^24^. Future research can explore whether different concentrations of sucralose interact with distinct taste receptors in both sweet and bitter GRNs.

Aspartame and Reb A are 200 and 400 times sweeter than sucrose in humans, respectively^6,80^. However, both sweeteners possess unique chemical structures that may be less compatible with fly taste receptors, leading to weaker activation compared to sucralose. Sweet and bitter GRN co-activation by Reb A is consistent with previous work demonstrating attraction to the sweetener in mice^81^, but also molecular evidence showing that it interacts with both sweet and bitter human taste receptors^82,83^. Interestingly, Reb A is considered less bitter for humans than the other popular stevia derivative, stevioside^84^. Future studies with these naturally derived sweeteners are necessary to fully understand the mechanisms of Reb A processing and how these compare to other steviol glycosides. Conversely, aspartame only activated bitter GRNs and elicited additional peaks of activity upon stimulus removal. These bitter OFF-responses have been observed with other aversive tastants^78,79^ and may indicate that more complex cellular kinetics contribute to aspartame processing. Aspartame leading to cellular responses only in bitter cells aligns with a study in mice that reported minimal attraction to this sweetener^85^, suggesting the “sweetness” associated with this compound is not conserved in either model organism. There is also evidence indicating that aspartame interacts with TRPV1 receptors that are expressed in taste cells^86^, however the mechanisms underlying chemosensation of this particular sweetener, and their apparent divergence in humans, remain unclear.

Combining our cellular results with our behavioral findings, we propose a combinatorial encoding mechanism for sucralose that is concentration dependent. Sweet GRNs respond to low concentrations to signal feeding attraction, whereas high concentrations co-activate bitter GRNs and signal feeding avoidance (Fig. 4G). This encoding pattern represents a common theme for the *Drosophila* peripheral gustatory system. Similar to sucralose, salts (NaCl)^48^, fatty acids (hexanoic acid)^55^, carboxylic acids (acetic acid)^54^, and amino acids (arginine)^56^ activate sweet GRNs at low concentrations to promote feeding while higher concentrations activate bitter GRNs to deter feeding. This appetitive-aversive balance likely enables flies to flexibly respond to chemicals that are beneficial to some extent, but harmful in high quantities. One caveat of our *in vivo* calcium imaging approach is that it does not directly measure neuronal activity, however previous investigation has confirmed that GCaMP signals are highly correlated with action potentials^87,88^. Future work using electrophysiological tip recordings can confirm if the level of action potentials induced in sweet and bitter GRNs by sweeteners is comparable to the calcium responses shown here.

### Artificial sweeteners differ in their behavioral valence and impact of bitter signaling

Many of the previous *Drosophila* investigations of sweeteners have focused on how chronic intake affects several phenotypes, including feeding^32,33,38–41^. However, these studies often do not quantify baseline interest in the sweeteners or assume that they elicit only attractive/sweet sensory signals. Additionally, sensory-induced physiological or metabolic changes also impact how dietary manipulations affect the animal^89^, but these interactions are often not addressed in this current area of research. Here, we characterized and compared the behavioral valencies of several artificial sweeteners. The limited attraction we observed towards many of the chemicals in our labellar PER experiments is consistent with previous tarsal PER data suggesting similarly mild response levels^42^. This study also reported that flies show positive preferential consumption of sucralose, aspartame, and stevioside^42^. Consistent with these results, we found that flies exhibited a positive feeding preference for both sucralose and Reb A in our 2-choice FLIC assay. In contrast, we found that the flies do not show a preference for aspartame. This discrepancy could be explained by FLIC quantifying behavioral interactions with different food sources^73^ instead of measuring ingestive preference in a dye-based assay^42^. Moreover, the lack of aspartame feeding preference we observed agrees with our calcium imaging results and previous mouse data^85^, further indicating that the palatability of this sweetener may be specific to humans.

Although bitter GRNs respond to sucralose, aspartame, and Reb A, our silencing experiments indicate that bitter cell activation only significantly modulates sucralose feeding. This result provides further evidence that sucralose evokes a stronger behavioral valence compared to the other sweeteners, and that bitter co-activation significantly modulates sucralose feeding.

Additionally, our behavioral evidence indicates that flies respond appetitively to high-concentration sucralose upon first encounter, but the accompanying bitter co-activation generates an aversive aftertaste signal that suppresses responses to subsequent stimuli, consistent with behavioral studies in mammals^26–28^. It is also worth emphasizing that we specifically focused on mechanisms of peripheral sucralose encoding and did not explore how these opposing gustatory signals are centrally processed to ultimately modify feeding output. Evidence from human neuroimaging studies demonstrate that sucralose and sucrose activate similar primary taste pathways, but sucralose produces significantly weaker responses in the insula and regions of the midbrain associated with feeding behavior^90^. Future analysis is required to determine how sucralose interacts with both sweet and bitter downstream circuitry.

In conclusion, our results indicate that flies exhibit diverse behavioral and cellular responses to various classes of artificial sweeteners. Future studies utilizing these sweeteners in *Drosophila* should anticipate sensory responses that are not exclusively sweet or appetitive. Moreover, future work with sucralose should expect concentration-dependent effects on feeding behavior. The chemosensory experience associated with sweetener consumption should be considered when evaluating the impact of diets containing artificial sweeteners on various phenotypes to better understand the unintended consequences of these chemicals.

## ACKNOWLEDGEMENTS

We thank the Bloomington Stock Center (BDSC) for fly stocks. Graphics were generated in BioRender.com. This work was supported by new lab startup funds from the Biology Department and College of Arts and Sciences at the University of Vermont and by the CNRs and the Atip-Avenir grant (CNRS Biology).

## AUTHOR CONTRIBUTIONS

Conceptualization, P.Y.M. and M.S..; methodology, P.Y.M. and M.S.; investigation, C.A., J.G., I.C., S.F., S.D., J.C., K.A., P.Y.M. and M.S.; writing - original draft, C.A., P.Y.M. and M.S.; writing - review & editing, C.A., J.G., I.C., S.F., S.D., J.C., K.A., P.Y.M. and M.S.; visualization, C.A. and M.S.; resources, P.Y.M. and M.S.; supervision, P.Y.M. and M.S.; funding acquisition, P.Y.M. and M.S.

## DECLARATION OF INTERESTS

The authors declare no competing interests.

## METHODS

### Flies

Experimental flies were kept on regular cornmeal food at 25°C in 60% relative humidity. Mated female flies between 2-10 days old were used in all experiments. The fly stocks used for experiments are listed in the Key Resource Table and genotypes are indicated in each figure. The Canton-Special (CS) strain was used as the wild-type *Drosophila melanogaster* strain.

### Chemicals

A full list of chemicals with source information can be found in the Key Resources Table. All- *trans*-retinal was made up in 100% EtOH, kept at -20°C, and diluted to a final concentration of 1 mM with EtOH of the same dilution given as a vehicle. Other test solution stocks were made as follows: sucrose: 1 M, sucralose: 0.1, 1, 10,100, 500 mM; Saccharin: 0.001, 0.01, 0.1, 1, 5, 10 mM; Cyclamate: 0, 1, 10, 50, 100, 200 mM; Aspartame: 0.001, 0.01, 0.1, 1, 5, 10 mM; Neotame: 0.001, 0.01, 0.1, 1, 5, 10 mM; Rebaudioside A: 0.001, 0.01, 0.1, 1, 5, 10 mM, Maltitol: 1, 10, 50, 100, 200 mM; Caffeine: 100mM.

### Proboscis extension response (PER)

For labellar PER, flies were placed inside a pipette tip cut to size so that only the head was exposed. Flies were then sealed into the tube with tape and then adhered to a glass slide with double-sided tape. Flies were allowed 1 to 2 hours to recover before testing began. Flies were stimulated with water on their labella, and allowed to drink until satiated. Each fly was then stimulated with increasing concentration of tastant, and proboscis extension responses to each tastant were recorded. Flies were provided with water between each tastant. All stimuli were delivered with a paper. For the sweet prolonged taste modulation experiment, flies were stimulated with water on their labella, and allowed to drink until satiated. Each fly was then stimulated with water, followed by 50 mM sucralose. 20 s or 5 min later flies were tested again for water or sucralose 50 mM. For the bitter prolonged taste modulation experiment, flies were stimulated with water on their labella, and allowed to drink until satiated. Each fly was then stimulated with water, followed by 500 mM sucralose. 20 s or 5 min later flies were tested again for 500 mM sucralose or water.

### Optogenetic PER

Flies were collected and put on standard cornmeal food supplemented with either ATR or vehicle for two days. Flies were then transferred to food deprivation vials containing 1% agar with either ATR or vehicle for one day prior to experiment. All vials were kept at 25°C and covered with foil to reduce light exposure. Flies were mounted the same way as for PER, but in a low luminosity environment. Flies were stimulated with water on their labella, and allowed to drink until satiated. Prior to the experiment, a green-light LED (3.5 mW) was turned ON, the flies were stimulated according to the bitter prolonged taste modulation experiment: each fly was stimulated with 500 mM sucralose. 20 s later flies were tested again for 500 mM sucralose (Fig. 3C). The same protocol was used with aspartame 0.1 mM and 10 mM (Fig. S2C); and with Rebaudioside A 0.1 mM and 10 mM (Fig. S3C).

### Calcium imaging

*In vivo* imaging of labellar GRN axon terminals was performed as previously described^48^. First, mated female flies were CO_2_ anesthetized and mounted into a specialized chamber in which their heads are secured with nail polish. Once secured, the fly’s labellum was manually extended and waxed into this position to ensure that the proboscis was unobstructed. Flies recovered in a humidity chamber for one hour. After recovery, cuticle above the sub-esophageal zone (the imaging area) was dissected to expose the brain, and Adult Hemolymph-Like (AHL) solution (108 mM NaCl, 5 mM KCl, 4 mM NaHCO3, 1 mM NaH2PO4, 5 mM HEPES, 15 mM ribose, 2mM Ca2+, 8.2mM Mg2+, pH 7.5) was continuously applied to the brain. The imaging area was cleared of obstructions by cutting and removing the respiratory tissues and part of the esophagus. Flies were imaged with a 3i Spinning disc Confocal station (Zeiss upright microscope, 2Kx2K 40 fps sCMOS camera, CSU-W1 T1 50 mm spinning disc). During image acquisition with a 40x water immersion objective, AHL was used as the immersive solution.

Once positioned for imaging, baseline fluorescence was recorded for 5 seconds before the fly’s labellum was stimulated with the tastant for 5 seconds, and the recording continued for a total of 17 seconds. The tastant was directly applied to the labellum via a micromanipulator and a capillary tube which fit directly over the fly’s labellum. Each fly was exposed to water (negative control), increasing concentrations of indicated sweeteners, and a positive control (1 M sucrose for Gr64f GRNs, 100 mM caffeine for Gr66a GRNs). The camera exposure was set to 100ms and 170 frames were captured at a rate of 9.5 Hz in Slidebook. Videos were opened in FIJI where a ROI was drawn around the projections and the fluorescence signal over time extracted. Ten consecutive points were chosen as a baseline and ΔF/F was calculated as a z-score ((F – mean baseline F)/standard deviation baseline F) for each timepoint. The three consecutive highest points during the 5 second stimulation were averaged as the peak ON response. Flies with no visible response to the positive controls were excluded

### FLIC

FLIC experiments were performed as previously described^66,73^. For optogenetic FLIC assays, flies were collected and put on standard cornmeal food supplemented with either all-*trans*-retinal or vehicle for two days. Flies were then transferred to food deprivation vials containing 1% agar with either all-*trans*-retinal or vehicle for one day prior to experiment. All vials were kept at 25°C and covered with foil to reduce light exposure. Flies were then individually transferred into behavioral feeding chambers via mouth pipette. Within each chamber, flies had access to two food sources (two-choice FLIC assay). Both options were connected to distinct capacitance sensors on *Drosophila* Feeding Monitors (DFMs) (Sable Systems) that recorded the number of interactions between the fly and each respective food source. Interaction with one side activated a green-light LED within that chamber (FLIC opto-lid, Sable systems), while the other side did not produce any light activation. For chronic silencing assays, flies were transferred to 1% agar food deprivation vials for 1 day prior to experiment and kept at 25°C. Flies were then loaded into similar two-choice feeding chambers with standard FLIC lids (Sable Systems) that lacked LED activation.

For both assays, interactions between a fly and each food source were recorded for 3 hours. A custom R script, based off the Pletcher Lab, was used to analyze this raw output (full code on GitHub, see Key Resources Table). Any interactions detected during the loading process were removed to avoid potential artifacts caused by loading flies into the chambers. Data were analyzed similarly to previous publications to quantify preference index, feeding interactions, feeding events, and feeding event duration^66,73^. Measurements were taken every 200ms, minimum signal threshold was set to 10, and feeding threshold was set to 20. For the optogenetic assays, green-light LED activation threshold was also set to 20 and the green light had an intensity of 1.5 mW. An interaction was recorded if the signal exceeded the feeding threshold. Preference index was calculated for each fly using: ((interactions on side A – interactions on side B) / total interactions). A feeding event was defined as 10 consecutive interactions above the minimum threshold, as long as one of these readings reached the feeding threshold. Sequential periods of continuous interactions were combined into one single event if the gaps of inactivity between them lasted less than 1 second. Feeding event duration was measured in seconds. To account for potential baseline differences between feeding chambers, flies of a particular condition or genotype were varied by position on the DFMs. Data from chambers that exhibited disrupted signal detection, including 0 values or exceedingly high values (>5000 interactions), were removed. Flies that failed to significantly interact with the food (<15 interactions) were also excluded. Lastly, we applied a ROUT outlier test for each feeding metric (preference index, interactions, events, and event duration) to remove any other significant outliers.

### Quantification and statistical analysis

All statistical tests were performed in GraphPad Prism 10 software. Specific tests are indicated in the figure legends along with the number of replicates, which were generally chosen based on variance and effect sizes in accordance with previously published literature using the same assays. Experimental or genotype controls were always run in parallel. Behavioral assays were repeated over multiple days and genetic crosses. Data are plotted as mean +/- SEM in bar graphs and line graphs. Asterisks indicate *p<.05, **p<.01, ***p<.001, ****p<.0001.

## Key Resources Table

| REAGENT or RESOURCE | SOURCE | IDENTIFIER |
| --- | --- | --- |
| **Chemicals, peptides, and recombinant proteins** |  |  |
| Ethanol | Pharmco | 111000200 |
| Sucrose | Sigma-Aldrich | S7903 |
| Sucralose | Sigma-Aldrich | 69293 |
| Aspartame | Sigma-Aldrich | W700655 |
| Rebaudioside-A | Sigma-Aldrich | 01432 |
| Neotame | Interchim | DW615 |
| Maltitol | Sigma-Aldrich | M8892 |
| Saccharin | Sigma-Aldrich | S6047 |
| Cyclamate | Sigma-Aldrich | C9131 |
| Caffeine | Sigma-Aldrich | C0750 |
| Agar | Sigma-Aldrich | A1296 |
| All-*trans*-Retinal | Sigma-Aldrich | R2500 |
| **Experimental models: Organisms/strains** |  |  |
| *D.melanogaster*: *w1118* | Wellgenetics (isogenic control for *IR94e* knock-in) | RRID: BDSC_3605 |
| *D.melanogaster*: *Canton-S* | Musso et al., 2017 |  |
| *D.melanogaster*: *Gr64f-Gal4* | Dahanukar et al., 2007 | Flybase: FBtp0057275 |
| *D.melanogaster*: *Gr66a-Gal4* | Wang et al., 2004 | RRID: BDSC_28801<br>Flybase: FBtp0014660 |
| *D.melanogaster*: *UAS-Kir2.1* | Baines et al., 2001 | Flybase: FBti0017552 |
| *D.melanogaster*: *UAS-GtACR1* | Bloomington Drosophila Stock Center | RRID: BDSC_92983<br>Flybase: FBti0202147 |
| *D.melanogaster*: *UAS-GCaMP6f* | Bloomington Drosophila Stock Center | RRID: BDSC_52869<br>Flybase: FBti0156888 |
| **Software and algorithms** |  |  |
| ImageJ | Schneider et al., 2012 | RRID: SCR_002285,<br>SCR_003070<br>https://imagej.nih.gov/ij |
| Slidebook 2024 | 3i (Intelligent Imaging Innovations) | RRID: SCR_014300<br>https://www.intelligent-imaging.com/slidebook |
| R Studio (4.3.2) | RStudio Team | RRID: SCR_000432<br>https://www.rstudio.com/ |
| Illustrator | Adobe | RRID: SCR_010279<br>https://www.adobe.com |
| Prism 10 | Graphpad | RRID: SCR_002798<br>https://www.graphpad.com/scientificsoftware/prism/ |
| BioRender | BioRender | RRID: SCR_018361<br>https://www.biorender.com |

---

**Figure S1:**
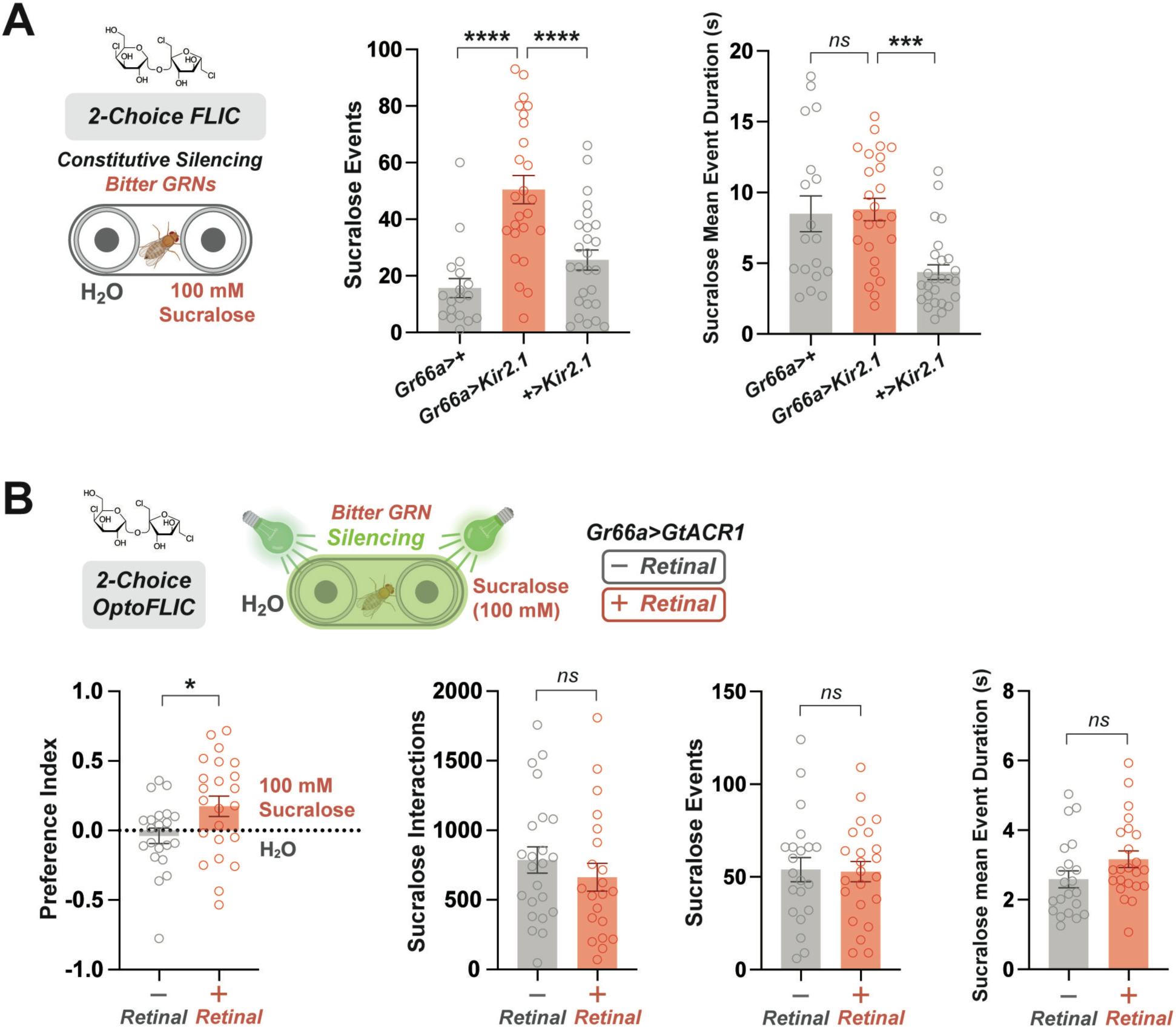
Constitutive and optogenetic silencing of bitter GRNs impacts sucralose feeding behavior. (A) Constitutive silencing of bitter GRNs (*Gr66a>Kir2.1*) during two-choice FLIC feeding assay. Flies were allowed to choose between two food sources, H_2_O and 100 mM sucralose for three hours. Number of feeding events and average feeding event duration for sucralose is depicted. *n* = 18-26 flies per genotype. (B) Optogenetic silencing of bitter GRNs (*Gr66a>GtACR1*) during two-choice OptoFLIC feeding assay. Flies pre-fed retinal (+Retinal) or control flies (-Retinal) were allowed to choose between two food options (H2O vs. 100 mM sucralose) for three hours. The green light LED was active for the duration of the experiment. Preference index calculated based off number of interactions with each food option. *n* = 18-23 flies per condition. All experiments used mated females. Data plotted as mean ± SEM. *ns =* no significance, *\*p < 0.05, ***p < 0.001, ****p < 0.0001* by ordinary one-way ANOVA with Dunnett’s multiple comparisons test (A) or unpaired t-test (B).

**Figure S2:**
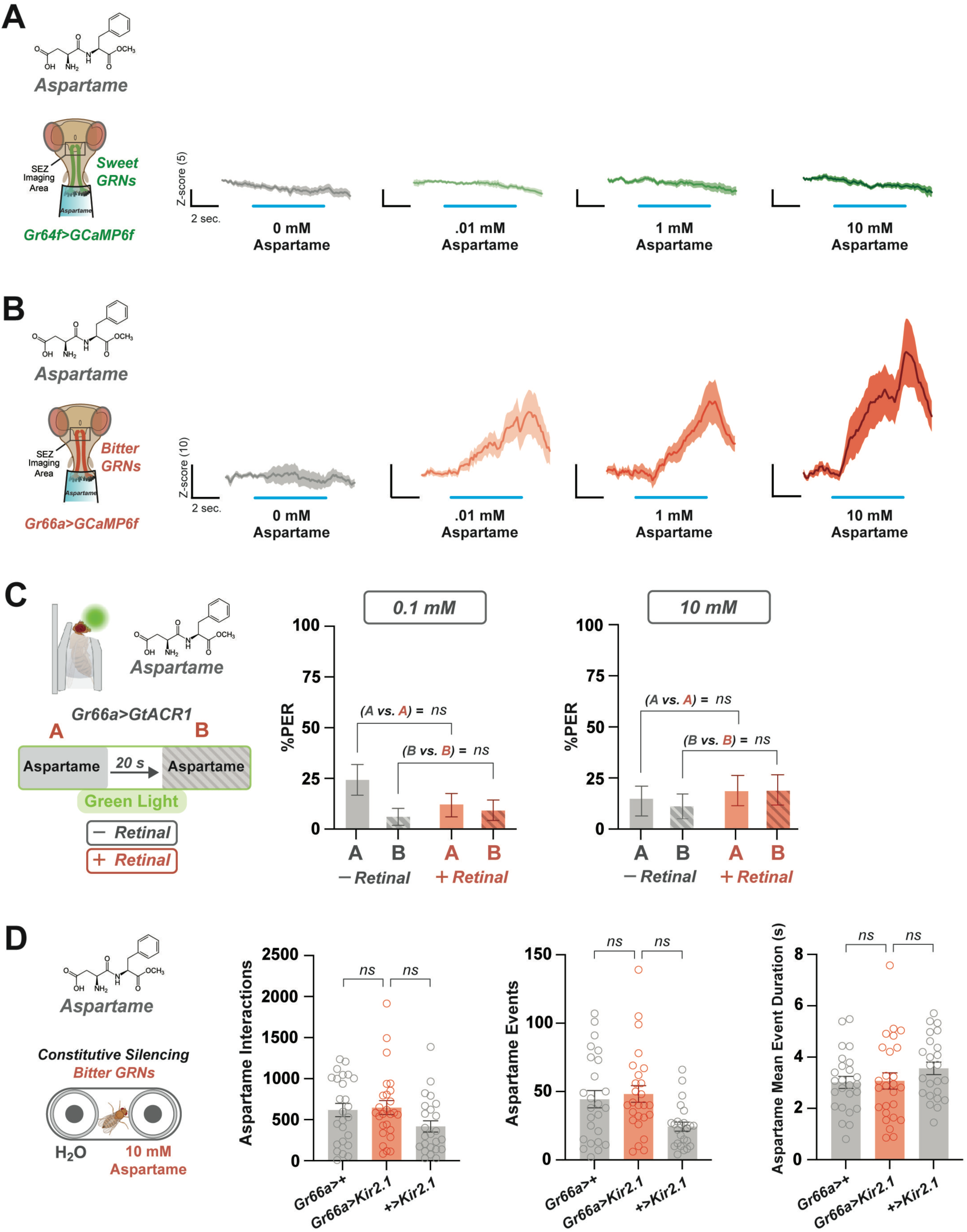
Aspartame elicits bitter activation but silencing bitter GRNs has no impact on feeding. (A,B) *In vivo* calcium imaging of sweet (A) and bitter (B) GRNs during labellar aspartame stimulation. Calcium responses measured as ΔF/F (Z-score) over time at each concentration. Blue lines under each curve indicate when the stimulus was on the labellum. *n* = 12 flies per experiment. (C) Optogenetic silencing of bitter GRNs (*Gr66a>GtACR1*) during prolonged taste modulation PER assay. Flies received two aspartame stimulations separated by 20 seconds while exposed to green light. Separate experiments were conducted at both a low (0.1 mM) and high (10 mM) aspartame concentration. %PER to the initial A and delayed B stimulations were compared between flies pre-fed retinal (+Retinal) and control flies (-Retinal). *n* = 25-34 flies per condition. (D) Constitutive silencing of bitter GRNs (*Gr66a>Kir2.1*) during two-choice FLIC feeding assay. Flies were allowed to choose between two food sources (H_2_O vs. 10 mM aspartame) for three hours. *n* = 26-27 flies per genotype. All experiments used mated females. All data plotted as mean ± SEM. *ns =* no significance by Wilcoxon matched-pairs signed rank test (C) or ordinary one-way ANOVA with Dunnett’s multiple comparisons test (D).

**Figure S3:**
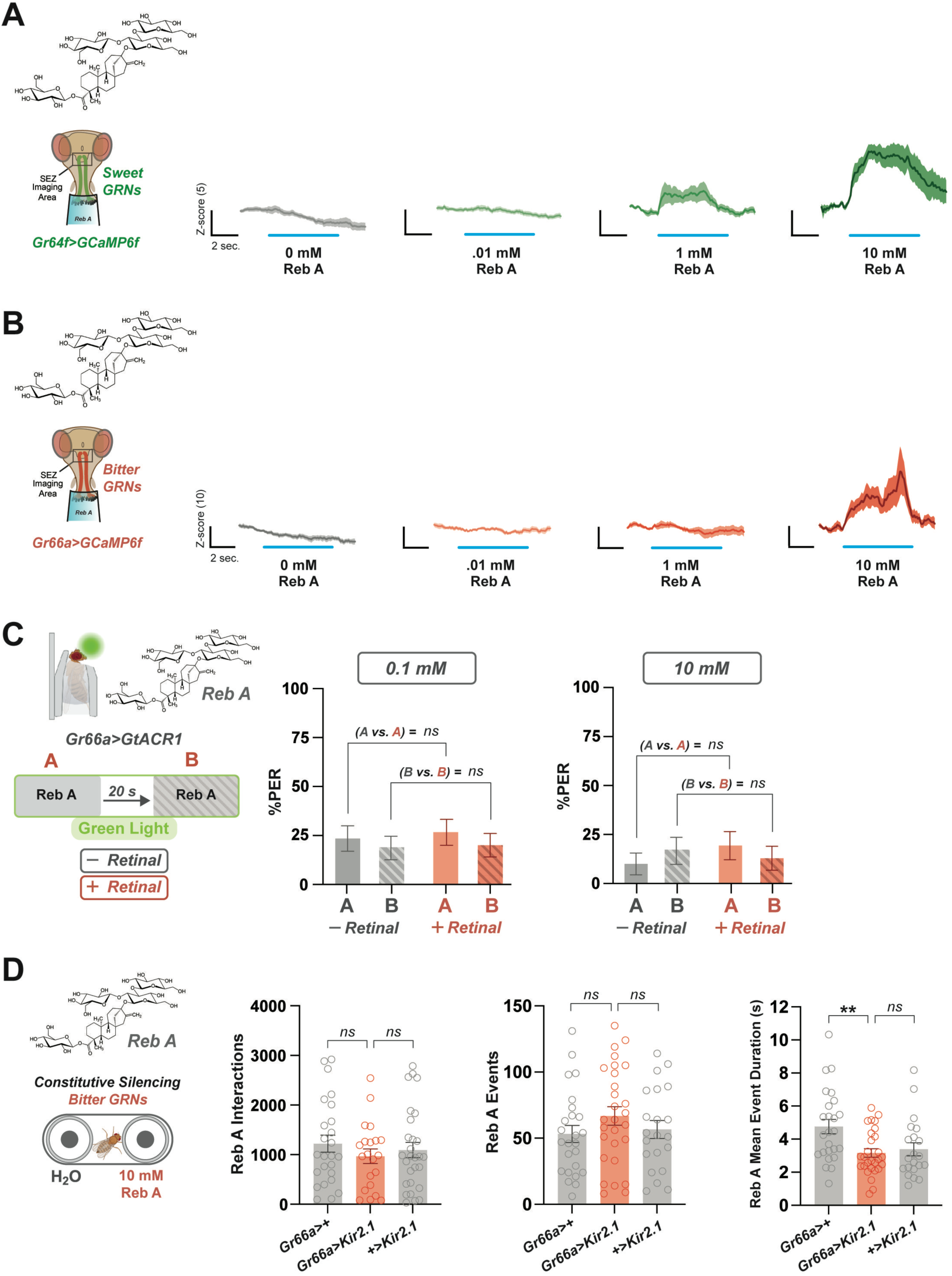
Reb A co-activates sweet and bitter GRNs but silencing bitter GRNs does not enhance feeding. (A,B) *In vivo* calcium imaging of sweet (A) and bitter (B) GRNs during labellar Reb A stimulation. Calcium responses measured as ΔF/F (Z-score) over time at each concentration. Blue lines under each curve indicate when the stimulus was on the labellum. *n* = 13 (A) and 11 (B). (C) Optogenetic silencing of bitter GRNs (*Gr66a>GtACR1*) during prolonged taste modulation PER assay. Flies received two Reb A stimulations separated by 20 seconds while exposed to green light. Separate experiments were conducted at both a low (0.1 mM) and high (10 mM) Reb A concentration. %PER to the initial A and delayed B stimulations were compared between flies pre-fed retinal (+Retinal) and control flies (-Retinal). *n* = 30-41 flies per condition. (D) Constitutive silencing of bitter GRNs (*Gr66a>Kir2.1*) during two-choice FLIC feeding assay. Flies were allowed to choose between two food sources (H2O vs. 10 mM Reb A) for three hours. *n* = 21-28 flies per genotype. All experiments used mated females. Data plotted as mean ± SEM. *ns =* no significance, *\*\*p < 0.01* by Wilcoxon matched-pairs signed rank test (C) or ordinary one-way ANOVA with Dunnett’s multiple comparisons test (D).

## Notes

### Competing Interest Statement

The authors have declared no competing interest.

## REFERENCES

1 Pearlman, M., Obert, J. & Casey, L. The Association Between Artificial Sweeteners and Obesity. Current Gastroenterology Reports 19, 64 (2017). 10.1007/s11894-017-0602-9

2 Sylvetsky, A. C. et al. Consumption of Low-Calorie Sweeteners among Children and Adults in the United States. Journal of the Academy of Nutrition and Dietetics 117, 441–448.e442 (2017). 10.1016/j.jand.2016.11.004

3 Sylvetsky, A. C. & Rother, K. I. Trends in the consumption of low-calorie sweeteners. Physiol Behav 164, 446–450 (2016). 10.1016/j.physbeh.2016.03.030

4 Ghusn, W., Naik, R. & Yibirin, M. The Impact of Artificial Sweeteners on Human Health and Cancer Association: A Comprehensive Clinical Review. Cureus 15, e51299 (2023). 10.7759/cureus.51299

5 Schiano, C. et al. Soft drinks and sweeteners intake: Possible contribution to the development of metabolic syndrome and cardiovascular diseases. Beneficial or detrimental action of alternative sweeteners? Food Research International 142, 110220 (2021). 10.1016/j.foodres.2021.110220

6 Peteliuk, V., Rybchuk, L., Bayliak, M., Storey, K. B. & Lushchak, O. Natural sweetener Stevia rebaudiana: Functionalities, health benefits and potential risks. Excli j 20, 1412–1430 (2021). 10.17179/excli2021-4211

7 Raben, A. & Richelsen, B. Artificial sweeteners: a place in the field of functional foods? Focus on obesity and related metabolic disorders. Curr Opin Clin Nutr Metab Care 15, 597–604 (2012). 10.1097/MCO.0b013e328359678a

8 Lohner, S., Toews, I. & Meerpohl, J. J. Health outcomes of non-nutritive sweeteners: analysis of the research landscape. Nutr J 16, 55 (2017). 10.1186/s12937-017-0278-x

9 Organization, W. H. Use of non-sugar sweeteners: WHO guideline. World Health Organization (2023). https://www.who.int/publications-detail-redirect/9789240073616

10 Ruiz-Ojeda, F. J., Plaza-Díaz, J., Sáez-Lara, M. J. & Gil, A. Effects of Sweeteners on the Gut Microbiota: A Review of Experimental Studies and Clinical Trials. Adv Nutr 10, S31–s48 (2019). 10.1093/advances/nmy037

11 Suez, J. et al. Artificial sweeteners induce glucose intolerance by altering the gut microbiota. Nature 514, 181–186 (2014). 10.1038/nature13793

12 Rodriguez-Palacios, A. et al. The Artificial Sweetener Splenda Promotes Gut Proteobacteria, Dysbiosis, and Myeloperoxidase Reactivity in Crohn’s Disease-Like Ileitis. Inflamm Bowel Dis 24, 1005–1020 (2018). 10.1093/ibd/izy060

13 Wang, X., Guo, J., Liu, Y., Yu, H. & Qin, X. Sucralose Increased Susceptibility to Colitis in Rats. Inflamm Bowel Dis 25, e3–e4 (2019). 10.1093/ibd/izy196

14 Swithers, S. E. Artificial sweeteners produce the counterintuitive effect of inducing metabolic derangements. Trends in Endocrinology & Metabolism 24, 431–441 (2013). 10.1016/j.tem.2013.05.005

15 Iizuka, K. Is the Use of Artificial Sweeteners Beneficial for Patients with Diabetes Mellitus? The Advantages and Disadvantages of Artificial Sweeteners. Nutrients 14, 4446 (2022).

16 Pepino, M. Y. Metabolic effects of non-nutritive sweeteners. Physiology & Behavior 152, 450–455 (2015). 10.1016/j.physbeh.2015.06.024

17 Witkowski, M. et al. The artificial sweetener erythritol and cardiovascular event risk. Nature Medicine 29, 710–718 (2023). 10.1038/s41591-023-02223-9

18 Debras, C. et al. Artificial sweeteners and risk of cardiovascular diseases: results from the prospective NutriNet-Santé cohort. BMJ 378, e071204 (2022). 10.1136/bmj-2022-071204

19 Chattopadhyay, S., Raychaudhuri, U. & Chakraborty, R. Artificial sweeteners – a review. Journal of Food Science and Technology 51, 611–621 (2014). 10.1007/s13197-011-0571-1

20 Xu, H. et al. Different functional roles of T1R subunits in the heteromeric taste receptors. Proc Natl Acad Sci U S A 101, 14258–14263 (2004). 10.1073/pnas.0404384101

21 Nie, Y., Vigues, S., Hobbs, J. R., Conn, G. L. & Munger, S. D. Distinct contributions of T1R2 and T1R3 taste receptor subunits to the detection of sweet stimuli. Curr Biol 15, 1948–1952 (2005). 10.1016/j.cub.2005.09.037

22 Moran, A. W. et al. Sweet taste receptor expression in ruminant intestine and its activation by artificial sweeteners to regulate glucose absorption. Journal of Dairy Science 97, 4955–4972 (2014). 10.3168/jds.2014-8004

23. 23 Brune, N. E. I. et al. (Google Patents, 2013).

24 Kuhn, C. et al. Bitter taste receptors for saccharin and acesulfame K. J Neurosci 24, 10260–10265 (2004). 10.1523/jneurosci.1225-04.2004

25 Lasconi, C. et al. Bitter tastants and artificial sweeteners activate a subset of epithelial cells in acute tissue slices of the rat trachea. Scientific Reports 9, 8834 (2019). 10.1038/s41598-019-45456-w

26 Roudnitzky, N. et al. Genomic, genetic and functional dissection of bitter taste responses to artificial sweeteners. Human Molecular Genetics 20, 3437–3449 (2011). 10.1093/hmg/ddr252

27 Torregrossa, A. M., Loney, G. C., Smith, J. C. & Eckel, L. A. Examination of the perception of sweet- and bitter-like taste qualities in sucralose preferring and avoiding rats. Physiol Behav 140, 96–103 (2015). 10.1016/j.physbeh.2014.12.023

28 Schiffman, S. S., Booth, B. J., Losee, M. L., Pecore, S. D. & Warwick, Z. S. Bitterness of sweeteners as a function of concentration. Brain Res Bull 36, 505–513 (1995). 10.1016/0361-9230(94)00225-p

29 Yarmolinsky, D. A., Zuker, C. S. & Ryba, N. J. Common sense about taste: from mammals to insects. Cell 139, 234–244 (2009). 10.1016/j.cell.2009.10.001

30 Tagorti, G., Yalçın, B., Güneş, M., Burgazlı, A. Y. & Kaya, B. Comparative evaluation of natural and artificial sweeteners from DNA damage, oxidative stress, apoptosis, to development using Drosophila melanogaster. Drug and Chemical Toxicology 47, 606–617 (2024). 10.1080/01480545.2023.2228522

31 Musso, P.-Y., Lampin-Saint-Amaux, A., Tchenio, P. & Preat, T. Ingestion of artificial sweeteners leads to caloric frustration memory in Drosophila. Nature Communications 8, 1803 (2017). 10.1038/s41467-017-01989-0

32 Hubrecht, I., Baenas, N., Sina, C. & Wagner, A. E. Effects of non-caloric artificial sweeteners on naïve and dextran sodium sulfate-exposed Drosophila melanogaster. Food Frontiers 3, 728–735 (2022). 10.1002/fft2.147

33 Adedayo, B. C., Akinniyi, S. T., Ogunsuyi, O. B. & Oboh, G. In the quest for the ideal sweetener: Aspartame exacerbates selected biomarkers in the fruit fly (Drosophila melanogaster) model of Alzheimer’s disease more than sucrose. Aging Brain 4, 100090 (2023). 10.1016/j.nbas.2023.100090

34 Vasconcelos, M. A., Orsolin, P. C., Silva-Oliveira, R. G., Nepomuceno, J. C. & Spanó, M. A. Assessment of the carcinogenic potential of high intense-sweeteners through the test for detection of epithelial tumor clones (warts) in Drosophila melanogaster. Food and Chemical Toxicology 101, 1–7 (2017). 10.1016/j.fct.2016.12.028

35 Umezaki, Y. et al. Taste triggers a homeostatic temperature control in hungry flies. eLife 13, RP94703 (2024). 10.7554/eLife.94703

36 Dus, M., Min, S., Keene, A. C., Lee, G. Y. & Suh, G. S. Taste-independent detection of the caloric content of sugar in Drosophila. Proc Natl Acad Sci U S A 108, 11644–11649 (2011). 10.1073/pnas.1017096108

37 Abu, F. et al. Communicating the nutritional value of sugar in *Drosophila*. Proceedings of the National Academy of Sciences 115, E2829–E2838 (2018). doi:10.1073/pnas.1719827115

38 Hasegawa, T. et al. Sweetness induces sleep through gustatory signalling independent of nutritional value in a starved fruit fly. Scientific Reports 7, 14355 (2017). 10.1038/s41598-017-14608-1

39 Wang, Q. P. et al. Sucralose Promotes Food Intake through NPY and a Neuronal Fasting Response. Cell Metab 24, 75–90 (2016). 10.1016/j.cmet.2016.06.010

40 Wang, Q.-P., Simpson, S. J., Herzog, H. & Neely, G. G. Chronic Sucralose or L-Glucose Ingestion Does Not Suppress Food Intake. Cell Metabolism 26, 279–280 (2017). 10.1016/j.cmet.2017.07.002

41 Park, J. H., Carvalho, G. B., Murphy, K. R., Ehrlich, M. R. & Ja, W. W. Sucralose Suppresses Food Intake. Cell Metab 25, 484–485 (2017). 10.1016/j.cmet.2017.02.011

42 Gordesky-Gold, B., Rivers, N., Ahmed, O. M. & Breslin, P. A. Drosophila melanogaster prefers compounds perceived sweet by humans. Chem Senses 33, 301–309 (2008). 10.1093/chemse/bjm088

43 Freeman, E. G. & Dahanukar, A. Molecular neurobiology of Drosophila taste. Curr Opin Neurobiol 34, 140–148 (2015). 10.1016/j.conb.2015.06.001

44 Singh, R. N. Neurobiology of the gustatory systems of Drosophila and some terrestrial insects. Microsc Res Tech 39, 547–563 (1997). 10.1002/(sici)1097-0029(19971215)39:6<547::Aid-jemt7>3.0.Co;2-a

45 Scott, K. Gustatory Processing in Drosophila melanogaster. Annu Rev Entomol 63, 15–30 (2018). 10.1146/annurev-ento-020117-043331

46 Fujii, S. et al. Drosophila sugar receptors in sweet taste perception, olfaction, and internal nutrient sensing. Curr Biol 25, 621–627 (2015). 10.1016/j.cub.2014.12.058

47 Jiao, Y., Moon, S. J., Wang, X., Ren, Q. & Montell, C. Gr64f is required in combination with other gustatory receptors for sugar detection in Drosophila. Curr Biol 18, 1797–1801 (2008). 10.1016/j.cub.2008.10.009

48 Jaeger, A. H. et al. A complex peripheral code for salt taste in Drosophila. Elife 7 (2018). 10.7554/eLife.37167

49 Marella, S. et al. Imaging Taste Responses in the Fly Brain Reveals a Functional Map of Taste Category and Behavior. Neuron 49, 285–295 (2006). 10.1016/j.neuron.2005.11.037

50 Deere, J. U. & Devineni, A. V. Taste cues elicit prolonged modulation of feeding behavior in Drosophila. iScience 25, 105159 (2022). 10.1016/j.isci.2022.105159

51 Ugur, B., Chen, K. & Bellen, H. J. Drosophila tools and assays for the study of human diseases. Dis Model Mech 9, 235–244 (2016). 10.1242/dmm.023762

52 Zhang, Y. et al. Fast and sensitive GCaMP calcium indicators for imaging neural populations. Nature 615, 884–891 (2023). 10.1038/s41586-023-05828-9

53 Stanley, M., Ghosh, B., Weiss, Z. F., Christiaanse, J. & Gordon, M. D. Mechanisms of lactic acid gustatory attraction in Drosophila. Curr Biol 31, 3525–3537.e3526 (2021). 10.1016/j.cub.2021.06.005

54 Devineni, A. V., Sun, B., Zhukovskaya, A. & Axel, R. Acetic acid activates distinct taste pathways in Drosophila to elicit opposing, state-dependent feeding responses. Elife 8 (2019). 10.7554/eLife.47677

55 Pradhan, R. N., Shrestha, B. & Lee, Y. Molecular Basis of Hexanoic Acid Taste in Drosophila melanogaster. Mol Cells 46, 451–460 (2023). 10.14348/molcells.2023.0035

56 Aryal, B., Dhakal, S., Shrestha, B. & Lee, Y. Molecular and neuronal mechanisms for amino acid taste perception in the Drosophila labellum. Curr Biol 32, 1376–1386.e1374 (2022). 10.1016/j.cub.2022.01.060

57 Arntsen, C., Guillemin, J., Audette, K. & Stanley, M. Tastant-receptor interactions: insights from the fruit fly. Frontiers in Nutrition 11 (2024). 10.3389/fnut.2024.1394697

58 Stocker, R. F. The organization of the chemosensory system in Drosophila melanogaster: a review. Cell Tissue Res 275, 3–26 (1994). 10.1007/bf00305372

59 Shiraiwa, T. & Carlson, J. R. Proboscis extension response (PER) assay in Drosophila. J Vis Exp, 193 (2007). 10.3791/193

60. 60 Das, A. & Chakraborty, R. in Encyclopedia of Food and Health (eds Benjamin Caballero, Paul M. Finglas, & Fidel Toldrá) 234-240 (Academic Press, 2016).

61 Dweck, H. K. M., Talross, G. J. S., Luo, Y., Ebrahim, S. A. M. & Carlson, J. R. Ir56b is an atypical ionotropic receptor that underlies appetitive salt response in Drosophila. Current Biology 32, 1776–1787.e1774 (2022). 10.1016/j.cub.2022.02.063

62 Saraiva, A., Carrascosa, C., Raheem, D., Ramos, F. & Raposo, A. Maltitol: Analytical Determination Methods, Applications in the Food Industry, Metabolism and Health Impacts. Int J Environ Res Public Health 17 (2020). 10.3390/ijerph17145227

63 Inagaki, Hidehiko K. et al. Visualizing Neuromodulation In Vivo: TANGO-Mapping of Dopamine Signaling Reveals Appetite Control of Sugar Sensing. Cell 148, 583–595 (2012). 10.1016/j.cell.2011.12.022

64 Wang, Q.-P. et al. PGC1α Controls Sucrose Taste Sensitization in Drosophila. Cell Reports 31, 107480 (2020). 10.1016/j.celrep.2020.03.044

65 Olivier, B. et al. Review of the nutritional benefits and risks related to intense sweeteners. Arch Public Health 73, 41 (2015). 10.1186/s13690-015-0092-x

66 Guillemin, J. et al. Taste cells expressing Ionotropic Receptor 94e reciprocally impact feeding and egg laying in Drosophila. Cell Rep 43, 114625 (2024). 10.1016/j.celrep.2024.114625

67 McDowell, S. A. T., Stanley, M. & Gordon, M. D. A molecular mechanism for high salt taste in Drosophila. Curr Biol 32, 3070–3081.e3075 (2022). 10.1016/j.cub.2022.06.012

68 May, C. E. et al. High Dietary Sugar Reshapes Sweet Taste to Promote Feeding Behavior in Drosophila melanogaster. Cell Rep 27, 1675–1685.e1677 (2019). 10.1016/j.celrep.2019.04.027

69 Shiu, P. K., Sterne, G. R., Engert, S., Dickson, B. J. & Scott, K. Taste quality and hunger interactions in a feeding sensorimotor circuit. Elife 11 (2022). 10.7554/eLife.79887

70 Govorunova, E. G., Sineshchekov, O. A., Janz, R., Liu, X. & Spudich, J. L. Natural light-gated anion channels: A family of microbial rhodopsins for advanced optogenetics. Science 349, 647–650 (2015). doi:10.1126/science.aaa7484

71 Musso, P.-Y. et al. Closed-loop optogenetic activation of peripheral or central neurons modulates feeding in freely moving Drosophila. eLife 8, e45636 (2019). 10.7554/eLife.45636

72 Deere, J. U. et al. Selective integration of diverse taste inputs within a single taste modality. Elife 12 (2023). 10.7554/eLife.84856

73 Ro, J., Harvanek, Z. M. & Pletcher, S. D. FLIC: High-Throughput, Continuous Analysis of Feeding Behaviors in Drosophila. PLOS ONE 9, e101107 (2014). 10.1371/journal.pone.0101107

74 Baines, R. A., Uhler, J. P., Thompson, A., Sweeney, S. T. & Bate, M. Altered Electrical Properties in *Drosophila* Neurons Developing without Synaptic Transmission. The Journal of Neuroscience 21, 1523–1531 (2001). 10.1523/jneurosci.21-05-01523.2001

75 Weaver, K. J. et al. Behavioral dissection of hunger states in Drosophila. Elife 12 (2023). 10.7554/eLife.84537

76 Tao, R. & Cho, S. Consumer-Based Sensory Characterization of Steviol Glycosides (Rebaudioside A, D, and M). Foods 9 (2020).

77 Shaher, S. A., Mihailescu, D. F. & Amuzescu, B. Aspartame Safety as a Food Sweetener and Related Health Hazards. Nutrients 15 (2023).

78 Devineni, A. V., Deere, J. U., Sun, B. & Axel, R. Individual bitter-sensing neurons in Drosophila exhibit both ON and OFF responses that influence synaptic plasticity. Curr Biol 31, 5533–5546.e5537 (2021). 10.1016/j.cub.2021.10.020

79 Snell, N. J. et al. Complex representation of taste quality by second-order gustatory neurons in Drosophila. Curr Biol 32, 3758–3772.e3754 (2022). 10.1016/j.cub.2022.07.048

80 Czarnecka, K. et al. Aspartame-True or False? Narrative Review of Safety Analysis of General Use in Products. Nutrients 13 (2021). 10.3390/nu13061957

81 Sclafani, A., Bahrani, M., Zukerman, S. & Ackroff, K. Stevia and saccharin preferences in rats and mice. Chem Senses 35, 433–443 (2010). 10.1093/chemse/bjq033

82 Hao, S. et al. Steviol rebaudiosides bind to four different sites of the human sweet taste receptor (T1R2/T1R3) complex explaining confusing experiments. Communications Chemistry 7, 236 (2024). 10.1038/s42004-024-01324-x

83 Allen, A. L., McGeary, J. E. & Hayes, J. E. Rebaudioside A and Rebaudioside D bitterness do not covary with Acesulfame K bitterness or polymorphisms in TAS2R9 and TAS2R31. Chemosens Percept 6 (2013). 10.1007/s12078-013-9149-9

84 Goyal, S. K., Samsher & Goyal, R. K. Stevia (Stevia rebaudiana) a bio-sweetener: a review. Int J Food Sci Nutr 61, 1–10 (2010). 10.3109/09637480903193049

85 Bachmanov, A. A., Tordoff, M. G. & Beauchamp, G. K. Sweetener preference of C57BL/6ByJ and 129P3/J mice. Chem Senses 26, 905–913 (2001). 10.1093/chemse/26.7.905

86 Riera, C. E., Vogel, H., Simon, S. A. & Coutre, J. l. Artificial sweeteners and salts producing a metallic taste sensation activate TRPV1 receptors. *American Journal of Physiology-Regulatory*, Integrative and Comparative Physiology 293, R626–R634 (2007). 10.1152/ajpregu.00286.2007

87 Ohkura, M. et al. Genetically Encoded Green Fluorescent Ca2+ Indicators with Improved Detectability for Neuronal Ca2+ Signals. PLOS ONE 7, e51286 (2012). 10.1371/journal.pone.0051286

88 Streit, A. K., Fan, Y. N., Masullo, L. & Baines, R. A. Calcium Imaging of Neuronal Activity in Drosophila Can Identify Anticonvulsive Compounds. PLOS ONE 11, e0148461 (2016). 10.1371/journal.pone.0148461

89 Yao, Z. & Scott, K. Serotonergic neurons translate taste detection into internal nutrient regulation. Neuron 110, 1036–1050.e1037 (2022). 10.1016/j.neuron.2021.12.028

90 Frank, G. K. et al. Sucrose activates human taste pathways differently from artificial sweetener. Neuroimage 39, 1559–1569 (2008). 10.1016/j.neuroimage.2007.10.061

